# Linking Motor Working Memory to Explicit and Implicit Motor Learning

**DOI:** 10.1101/2025.02.27.640657

**Authors:** Hanna Hillman, Taylor McClure, Samuel D. McDougle

## Abstract

While explicit and implicit motor learning have been widely studied, the extent to which working memory for recent movements – motor working memory (MWM) – contributes to these processes remains unclear. Previous research suggests that visuospatial working memory may facilitate explicit motor learning but is either uninvolved in or detrimental to implicit learning. Here, we ask if and how these findings extend to non-visual MWM. Based on recent work pointing to separate effector-independent and effector-specific MWM codes, we hypothesized that: (1) explicit motor learning processes would correlate with effector-independent MWM, and (2) implicit motor learning processes would correlate with effector-specific MWM. To test these hypotheses, human participants completed both a MWM task and a visuomotor adaptation task. Our results revealed significant correlations between the quality of effector-independent MWM and the degree of explicit motor learning, extending previous findings on visuospatial working memory. Additionally, we present evidence supporting our second hypothesis, that effector-specific MWM correlates with implicit motor learning.

## Introduction

Working memory and skill learning have a dynamic relationship. Consider learning a new dance move: with each less-than-perfect attempt, you can use remembered details of what went right or wrong in the previous attempt to guide your next try. To complicate matters, substantial research suggests that working memory—the process of actively maintaining and manipulating information over short timescales—is not a unitary process. Instead, it involves specialized functional and anatomical resources that vary depending on the type of information being processed (E. J. Adams et al., 2018; Levy & Goldman-Rakic, 2000). Visual, auditory, and verbal working memory are widely studied in cognitive psychology and neuroscience. However, how working memory processes movement-related information—also known as Motor Working Memory (MWM)—and its relationship with motor learning remains unclear.

What do we know about MWM? Classic work examining short-term recall of arm positions without visual feedback have revealed archetypal working memory effects, including set size effects, interference effects, and rapid forgetting (J. A. Adams & Dijkstra, 1966; Posner & Konick, 1966; Schmidt et al., 2018; Schmidt & Ascoli, 1970, 1970; Smyth et al., 1988; Smyth & Pendleton, 1989). Moreover, neurophysiology in non-human primates has revealed retrospective short-term memory traces of recent eye movement kinematics, manifested as persistent neural firing in the dorsolateral prefrontal cortex (Calangiu et al., 2025; Tsujimoto & Sawaguchi, 2004).

Similar to how visual working memory differentiates between spatial (“where”) and object-based (“what”) features, recent work from our lab suggests that MWM represents movement information in at least two ways: effector-specific and effector-independent (Hillman et al., 2024). Effector-specific information refers to content – such as proprioceptive information – that is strictly associated with the exact body parts involved in an action. On the other hand (so to speak), effector-independent information includes abstract content that can be transferred across effectors or even across individuals (e.g., during imitation; Wong et al., 2019). There is evidence that effector-independent information may be maintained without interference from a simultaneous load in visuospatial working memory, suggesting that MWM may constitute its own distinct working memory subsystem rather than being reducible to visuospatial and proprioceptive memory (Hillman et al., 2024). Returning to our dance example, if your new dance move involves waving your right arm in a circular motion, you may encode both effector-specific information – such as the kinesthetic sensation of moving your arm in that manner – and effector-independent information, such as the spatial trajectory your hand traced through space.

Motor learning can also be divided into distinct processes. The most prominent distinction is between explicit motor learning, which involves deliberate decision-making and allows for the intentional application of cognitive movement strategies, and implicit motor learning, which involves slow, unconscious recalibration that gradually alters internal sensorimotor mappings (Krakauer et al., 2019; McDougle et al., 2015; McDougle, Ivry, et al., 2016; McDougle et al., 2017; Schween et al., 2020; Taylor et al., 2014). Early motor learning is largely characterized by explicit processes, during which the learner relies on time-consuming conscious strategies that constrain the range of possible actions and adjust for gross errors (Albert et al., 2022; Fitts & Posner, 1967; Krakauer et al., 2019; Langsdorf et al., 2021; Tsay et al., 2024). Over time, as movements become more automatic and efficient, implicit processes become more dominant. The dissociation between explicit and implicit motor learning is often studied using visuomotor adaptation tasks, in which participants have to compensate for a perturbed (typically rotated) mapping between sensory feedback (e.g., a cursor on a screen) and motor commands (e.g., reaching movements) (Cunningham, 1989; Krakauer, 2009). Explicit learning in these tasks primarily involves deliberately “re-aiming” movements to counteract performance errors, which requires remembering successful actions and often relies on spatial reasoning about the perturbed environment (McDougle & Taylor, 2019; Tsay et al., 2024). In contrast, implicit learning involves the fine-tuning of movement kinematics to gradually and unconsciously counteract sensory prediction errors (Taylor et al., 2014).

Prior research suggests that explicit motor learning relies on working memory, whereas implicit learning does not (Bo & Seidler, 2009; Christou et al., 2016; Masters, 1992; McNay & Willingham, 1998). Using fMRI, Anguera et al. (2010) observed activity in cortical areas associated with spatial working memory during the early phases of a visuomotor adaptation task—when explicit motor learning is most prevalent—but found limited shared activity in later learning, which primarily reflects implicit processes. Moreover, evidence suggests that working memory is not merely uninvolved in later implicit learning but may actively disrupt it. For example, top-down conscious control processes, such as working memory, can impair the automation of motor skills (Masters, 1992). Christou et al., (2016) found a positive relationship between spatial working memory capacity and explicit learning in a visuomotor adaptation task, as well as a negative relationship between spatial working memory and implicit learning. To date, research on the relationship between working memory and motor learning has primarily focused on visual working memory. However, it remains unclear whether and how these findings generalize to motor working memory (MWM).

Here, we investigated whether the dissociable components of MWM are differentially linked to explicit and implicit motor learning processes. Specifically, we hypothesized that: (1) Effector-independent MWM ability would be selectively correlated with the degree of explicit motor learning, supporting the idea that explicit motor learning is linked not only to visuospatial working memory but also to working memory processes specialized for the short-term storage of abstract motor commands. (2) Effector-specific MWM would correlate positively with implicit motor learning. The second prediction is more speculative, as it challenges the notion that working memory is either independent of or disruptive to implicit motor learning. If confirmed, this would suggest that it is not working memory recruitment itself that impairs implicit motor learning but rather the specific content being maintained in working memory. To test these hypotheses, we designed a correlational study using two tasks: a visuomotor rotation (VMR) learning task that dissociated explicit and implicit motor learning and a non-visual MWM task that dissociated effector-specific and effector-independent components.

## Materials & Methods

### Participants

The study was reviewed and approved through Yale University’s Institutional Review Board, and all participants provided written informed consent prior to participation. A total of 34 participants were recruited through a psychology pool and received class credit for their participation. Two participants were excluded due to incomplete data collection, and one participant was excluded for aberrant explicit learning behavior (i.e., negative explicit learning in the VMR task, which was also less than three standard deviations from the mean explicit learning observed across the sample). Of the remaining 31 participants, 17 were female, with a mean age of 18.8 ± 0.75 years (SD), and 27 participants were right-handed, as assessed by the Edinburgh Handedness Inventory (Oldfield, 1971).

### Apparatus

Both tasks were conducted using a Kinarm End-Point Lab (BKIN Technologies Ltd., Kingston, ON, Canada), equipped with a low-friction, two-joint robotic arm fitted with a cylindrical handle that enabled planar reaching movements by passively guiding participants or being actively guided by them. Participants were seated in an adjustable chair with their feet on the ground in two outlined boxes, discouraging them from moving their legs throughout the experiment. The hand not holding the robot’s handle at any given moment rested flat on a mousepad on the workspace table. Participants sat with their foreheads resting against a soft pad mounted to a horizontal LCD monitor. The monitor was mounted parallel to the table facing a semi-silvered mirror, allowing them to comfortably view the presented instructions and visual stimuli while preventing them from seeing their hands below the mirror. Furthermore, participants wore a fabric bib around their necks, which was fastened to the mirror and fully occluded their upper bodies. An experimenter remained in the room for the duration of the study to monitor participants’ adherence to instructions and postural requirements.

### Motor Working Memory Task

The only visual stimuli participants received throughout the MWM task were text prompts that appeared outside the reaching workspace. At no point did participants see their hands, arms, or any indication of their arm position, nor did they receive any performance feedback throughout the task. The MWM task was always completed before the VMR task. Participants first completed a five-trial practice block (excluded from analysis) to become accustomed to the robot and ask any task-related questions. Once the experimenter was confident that the participant understood the instructions, they proceeded to complete two experimental blocks of 48 trials each. Participants were allowed to rest briefly between blocks but remained seated throughout.

Each trial consisted of three phases: encoding, maintenance, and retrieval (Figure 1A). Before each trial, participants were instructed to grasp the robot’s handle with their right hand, which then guided them to the “home” position, located approximately 13 cm from their chest along the midline. During the encoding phase, participants were passively guided by the robot in outward reaching trajectories to four different locations, returning to the home position between each movement. While moving, a number was displayed above the workspace, indicating the current movement in the sequence (e.g., participants saw “#1” during the first movement, “#2” during the second, and so on). Each movement lasted 800 ms. At the end of each outward movement, the robot paused for 1 s before returning to the home position, where the hand rested for an additional 1 s between movements. There were 12 possible invisible targets 9 cm from the home position, evenly spaced between 15° and 165°. Target locations were pseudorandomly selected to ensure equal sampling throughout the task, with no repetitions within a trial and a minimum angular difference of ±25° between consecutive movements in a given sequence.

**Figure 1.**
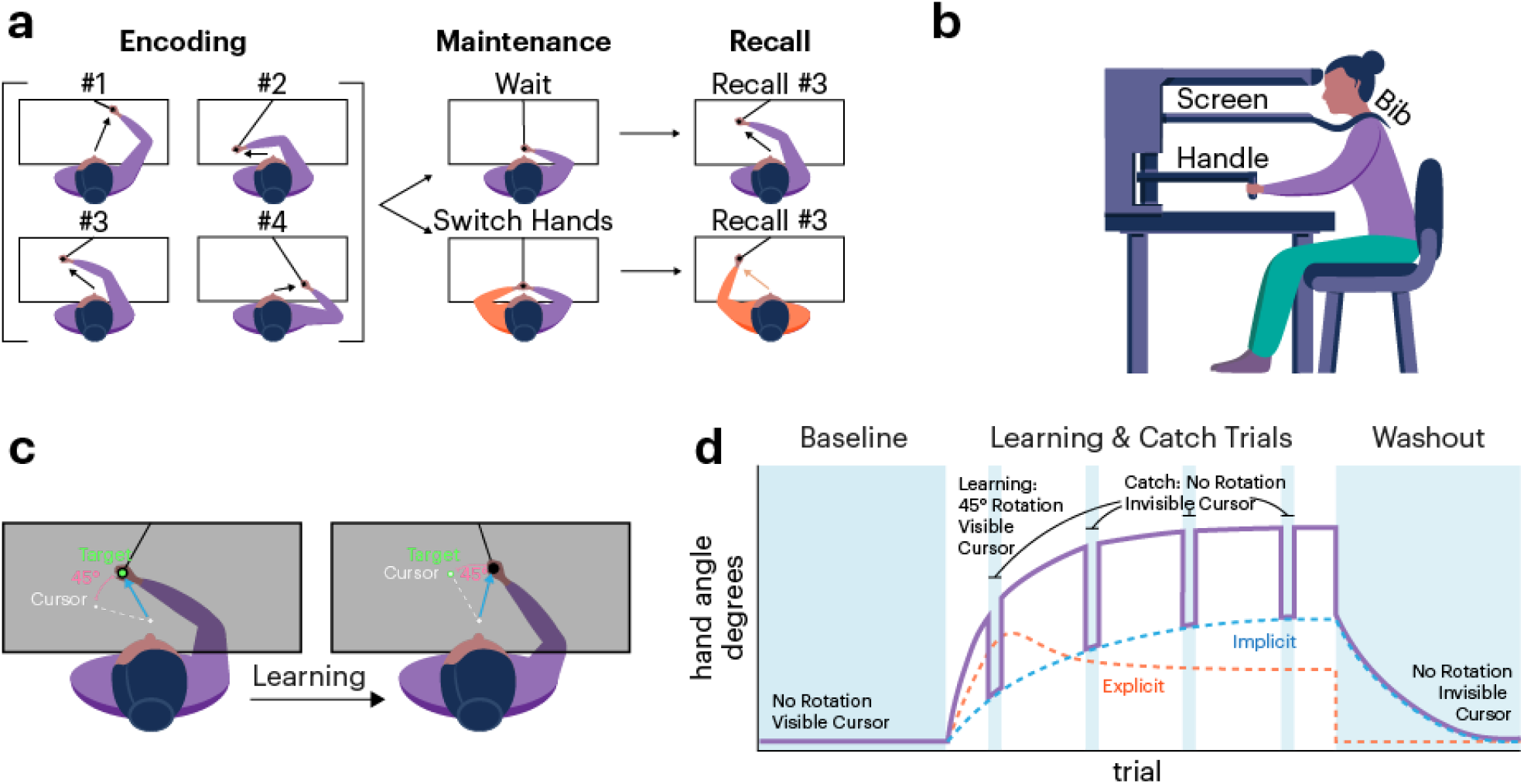
Motor working memory and visuomotor learning task design. (**a**) *Experimental setup used in both tasks*: Participants were seated at a table with their heads resting on a headrest and their eyes directed toward a screen that displayed visual stimuli while blocking any view of their hands and arms. The rest of the upper body was occluded by an opaque bib. (**b**) *MWM task*: During the encoding phase, participants were passively guided to four locations by the robotic arm, returning to a home position between movements. During each movement they saw a number corresponding to the current movement’s position in the sequence order. While maintaining this information in memory, participants were instructed either to wait or to switch the hand grasping the handle. Finally, they were prompted to recall one of the movements using whichever hand was currently on the handle. (**c**) *Visuomotor Rotation (VMR) task*: During the learning phase, participants were instructed to make rapid, straight movements to direct a cursor through a target. In rotation trials, the cursor was rotated 45° relative to their movement direction. Participants adjusted their movements to compensate for the cursor rotation. (**d**) *Schedule of trial types in the VMR task and multiple learning processes*: The VMR task consisted of four trial types. Participants first completed a baseline phase, during which they moved toward the targets while receiving veridical cursor feedback on their hand position. In the subsequent learning phase, the cursor was rotated relative to the hand position (as described in panel c), requiring participants to adapt their movements. During the learning phase, participants experienced four catch-trial periods in which they were instructed to move directly toward the target without compensating for rotation; during these trials, no cursor feedback was provided. This method isolated implicit motor adaptation, allowing for the dissociation of explicit and implicit contributions to learning (see *Methods*). Finally, in the washout phase, participants performed trials identical to the catch trials, allowing for the measurement of the final implicit learning state.

After encoding the four movements, participants were required to maintain all four movements in working memory for 3 s. During this period, they were instructed to either (1) “Switch hands,” in which case they would grasp the handle with their opposite (left) hand, or (2) “Wait,” keeping their right hand on the handle without letting go. Following the 3-second maintenance phase, participants were prompted to recall one of the four movements (e.g., “Recall movement #1”) using whichever hand was currently grasping the handle. To recall a movement, they executed an outward-reaching motion that matched their memory of the cued movement trajectory in the workspace (non-mirrored), then paused at the recalled movement’s endpoint. The robot registered their final location after 500 ms of dwell time before guiding them back to the home position. Participants were not informed in advance which movement they would be asked to recall or whether they would need to switch hands during maintenance. The specific movement probed for recall (i.e., the first, second, third, or fourth movement from the encoding sequence) was randomly selected and equally sampled.

A previous study from our lab demonstrated that this hand-switching paradigm is an effective method for dissociating effector-specific and effector-independent contributions to motor working memory (Hillman et al., 2024). When participants were instructed to recall a movement from working memory using the same hand they had used to encode it (the “Same” condition), they could, in theory, utilize both effector-specific (e.g., somatosensory) and effector-independent (e.g., abstract spatial trajectory) information to inform recall. However, when participants switched hands (the “Switch” condition) and recalled movements with the opposite hand, effector-specific information was no longer applicable, meaning they could rely only on effector-independent information to inform recall (Figure 1A)

### Visuomotor Rotation Task

Visuomotor rotation tasks are widely used to assess error-based motor learning and to measure the simultaneous contributions of implicit and explicit motor learning processes (Taylor et al., 2014). Participants attempted to move a small white visual cursor, representing their hand position (0.15 cm radius), through a green target (0.5 cm radius) (Figure 1C). A circular home position landmark (0.5 cm), the target, and, on most trials, the cursor were the only visual feedback provided during the task. As in the MWM task, participants could not see their hands or the robotic arm. On each trial, the target was positioned at either 0°, 60°, 120°, or 180°, located 9 cm from the central home position. Participants saw one target per trial, and target locations were randomly selected and equally sampled throughout the task. For each movement, participants were instructed to reach through the target with a fast, straight, ballistic movement. At the end of the movement, the robot guided them back to the home position. If participants waited more than 500 ms after the target appeared, they were instructed to initiate their movement sooner in the next trial. If their movement took more than 500 ms to cross the target radius, they were asked to move faster.

The task consisted of four types of blocks: baseline, rotation, catch, and washout. A schedule of these trials is shown in Figure 1D. The experiment began with 39 baseline trials (the 40th baseline trial was not recorded due to a software error). During baseline trials, participants received continuous online cursor feedback as they reached through a target. Following the baseline block, the rotation block commenced. In rotation trials, the cursor feedback was rotated ±45° relative to the reach direction, with the rotation direction counterbalanced across participants. Participants thus had to learn to counteract this 45° discrepancy to restore accurate performance. During the rotation blocks, participants received online feedback on their cursor position. The rotation phase consisted of five 32-trial blocks, totaling 160 rotation trials.

Crucially, we also included four brief “catch” blocks interspersed throughout the rotation phase. Each catch block consisted of four trials—one for each target location—presented in a random order. Before each catch block, participants were instructed to abandon any strategy they had been using to move the cursor to the target in the rotation trials, and instead reach directly toward the target. During these trials, participants did not see their cursor. Without error feedback and without the use of any explicit aiming strategy, the catch blocks served as a measure of participants’ implicit learning. Explicit learning was then inferred by subtracting movement angles in catch blocks from those in neighboring rotation trials. This approach, known as the exclusion method, has been shown to be an effective and straightforward technique for dissociating explicit and implicit motor learning (Maresch et al., 2020).

Finally, the experiment concluded with a 40-trial washout block. The instructions and feedback for these trials were identical to those of the catch trials. This block measured the final state of implicit learning and its gradual decay.

### Analysis

Data preprocessing was performed using MATLAB (MathWorks, 2024), and all analyses were conducted in R (R Core Team, 2024). For statistical tests, all reported t-tests are two-tailed, paired tests with an alpha level of 0.05. In our key correlational analyses, we report both Pearson and Spearman correlation coefficients (for completeness), also using an alpha level of 0.05.

For the MWM task, participant performance was assessed using two metrics: variability in memory errors and absolute memory errors, both in the angular and extent dimensions (see *Results*). Variability was quantified using the interquartile range (IQR) of angular errors, chosen for its robustness to outliers given the limited number of trials in some data subsets (see *Results*). This approach follows previous studies on both hand localization and working memory variability (Shin et al., 2017; Will & Stenner, 2024). Because our study was primarily correlational, we included multiple MWM metrics to evaluate the robustness of any observed correlations.

For the VMR task, behavior was quantified by computing participants’ hand angles relative to the start position at the moment they crossed the invisible ring defined by the start position (the ring’s center) and the target location. Three key VMR metrics were calculated: (1) Total learning, defined as the mean hand angle for the last three trials at each target location in the final rotation block (12 trials total); (2) Explicit learning, indirectly measured by subtracting the last catch block (a measure of implicit learning) from total learning; and (3)Total implicit learning (early aftereffects), calculated as the average hand angle for the first three trials at each target location in the washout block.

In addition to the three participants excluded (see *Participants*), individual trials were removed from analysis in each task based on performance. In the MWM task, 98 trials (3.3% of all data) were excluded for having an angular error over ±90°, suggestive of a memory lapse, hand slip, or swap error. In the VMR task, 51 trials (0.6% of total data) were removed for having a hand angle exceeding ±3 standard deviations from the block’s mean hand angle.

## Results

### Motor Working Memory Task

Participants (n = 31) in our study performed two tasks: a motor working memory (MWM) task and a visuomotor rotation adaptation task (Figure 1). Building on our recent finding (Hillman et al., 2024) that MWM is composed of two distinct representational codes – effector-independent and effector-specific – we hypothesized that these codes would correlate, respectively, with two dissociable components of visuomotor learning: explicit strategic learning and implicit motor adaptation. Such a finding would provide evidence for a link between well-known mechanisms of long-term motor learning and short-term motor memory processes.

To dissociate effector-specific and effector-independent contributions to MWM, participants passively encoded four movements and were then asked to recall one of them using either the same hand (Same condition) or the opposite hand (Switch condition; see Methods and Figure 1A). We reasoned that in the Same condition, both effector-specific and effector-independent information could contribute to recall performance. However, in the Switch condition, where participants switched effectors (hands) between encoding and recall, only effector-independent information could influence recall performance.

The results of the MWM experiment replicated our previous finding that the benefit of using the same hand for encoding and recall is limited to the most recently encoded movements (Figure 2; Hillman et al., 2024). Two metrics were used to quantify MWM performance, angular variability (IQR) in report errors and absolute angular errors (see *Methods*). For IQR, a significant difference between Switch and Same errors was observed when the fourth (last-encoded) movement was cued for recall (t(29) = 2.28, p = 0.029, d = 0.41). However, no significant differences were found when the first (t(29) = 0.84, p = 0.409, d = 0.15), second (t(29) = 0.56, p = 0.579, d = 0.10), or third (t(29) = 1.46, p = 0.155, d = 0.26) movements were recalled. For the mean of absolute angular errors a significant difference between Switch and Same errors was observed for the fourth (t(29) = 4.62, p = 0.0001, d = 0.83) and third (t(29) = 2.21, p = 0.035, d = 0.40) movements, but not for the first (t(29) = 1.34, p = 0.191, d = 0.24) or second (t(29) = 1.26, p = 0.216, d = 0.23) movements.

**Figure 2.**
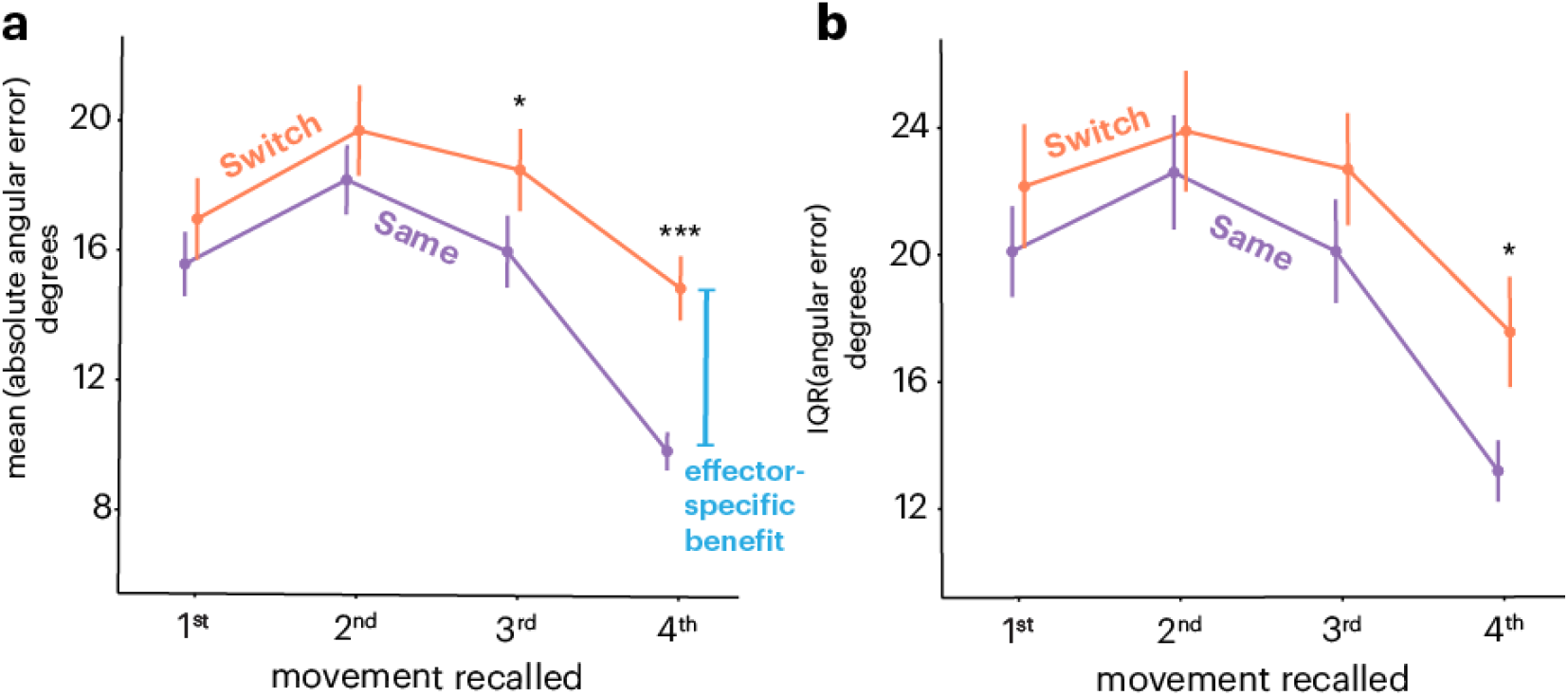
MWM results. Movements on the x-axis are ordered by sequence position, where “1st” represents the first movement encoded (and thus the oldest). The y-axis reflects participants’ errors in each condition, measured as either (a)the average absolute angular error or (**b**) variability, as the interquartile range (IQR) of angular errors. The difference between these conditions indicates the advantage of using the same hand for recall, referred to as the effector-specific benefit. Error bars represent the standard error of the mean (s.e.m.). *p < .05; ***p < .005.

These results align with our recent findings, which demonstrate that working memory for recently encoded movements exhibits attenuated interlimb transfer, whereas movements encoded earlier undergo successful transfer (Hillman et al., 2024). Our previous work attributed this pattern to interference at the encoding limb rather than passive temporal decay. Specifically, we found evidence that as additional movements are made, they interfere retroactively with the effector-specific (but not effector-independent) memory (Figure 2).

Effector-independent information is available in all trial conditions (Switch and Same); therefore, isolating effector-specific contributions to performance is more complicated. To approximate effector-specific MWM, two methods were used: (1) an “effector-specific benefit” was calculated by subtracting the Same condition’s performance error from that of the Switch condition (note that subtracting the Switch from Same would not calculate benefit, as the comparison is based on error). This difference was measured only for the fourth movement, as it was the sole position that showed a reliable difference between the Same and Switch conditions across both MWM metrics. (2) A second, sequence-dependent “fourth-movement” metric was quantified by calculating the difference between performance error in the fourth position and that of the other three positions within the Same condition. As previously mentioned, evidence suggests that as new movements are executed, they retroactively interfere with or “wash out” previously stored effector-specific information (Hillman et al., 2024). Therefore, by comparing the fourth movement, which shows significant effector-specific benefit, with the other three movements, effector-specific information can be inferred. Since this measurement is derived only from the Same condition, it has the added advantage of not being subject to performance disruptions that could arise from the act of switching hands during maintenance.

All three of these variables—effector-independent MWM and the two estimates of effector-specific MWM—were then correlated with key measures from the visuomotor rotation learning task (implicit and explicit) to examine potential relationships between MWM codes and error-based motor learning. We revisit this analysis in our cross-task correlation section.

### Visuomotor Rotation Task

The same participants who performed our MWM task also completed a standard visuomotor rotation (VMR) task. Participants readily adapted to the 45° visuomotor rotation (Figure 3A), reaching an asymptotic hand angle of 36.21° ± 5.51° (*t*(29) = 36.63, *p* < 2.2e-16, *d* = 6.58).

**Figure 3.**
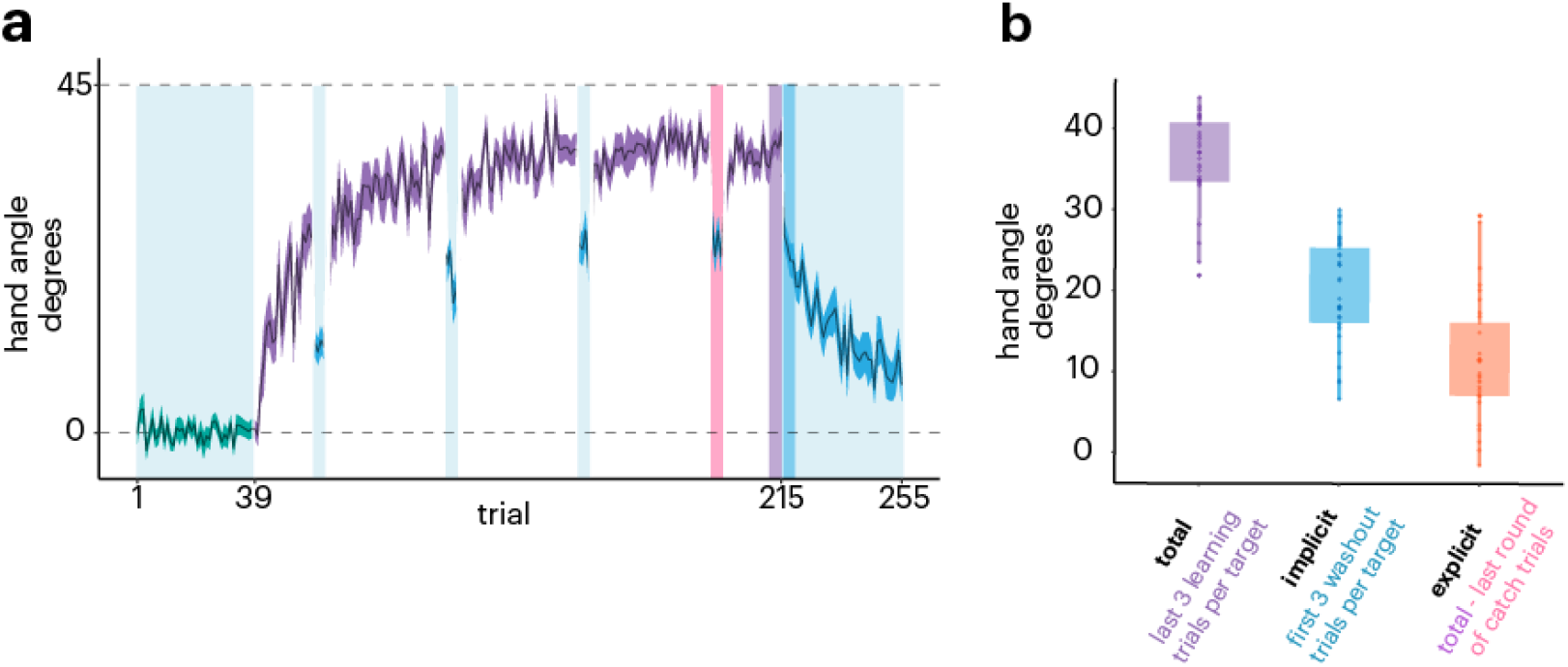
VMR results. (**a**) The mean hand angle per trial for each experimental condition (green = baseline; purple = learning phase; blue = catch trials and washout phase). Error shading reflects the s.e.m. (**b**) Total learning was calculated as the average of the last three learning trials per target (12 trials total). Implicit learning was measured using the first three washout trials per target. Explicit learning was determined by subtracting the final round of catch trials from the total learning metric. Box plot height shows confidence intervals and dots show individual subjects.

As described in the *Methods*, the “exclusion” method was used to isolate implicit learning by instructing participants to abandon any cognitive strategy, reach directly to the displayed targets, and receive no visual feedback on a subset of trials. This approach allowed explicit learning to be quantified through simple subtraction by deducting implicit learning from the measured hand angles on neighboring standard non-exclusion trials.

The dissociation of explicit and implicit learning was successful (Figure 3B), demonstrating robust contributions of both components to the overall learning curve. Asymptotic explicit learning reached 11.51° ± 7.52° (t(29) = 8.51, p < 1.7e-9, d = 1.53), while asymptotic implicit learning, measured over the first three cycles (12 trials) of the washout phase (early aftereffects), reached 19.83° ± 6.36° (t(29) = 17.37, p < 2.2e-16, d = 3.12). These two measures of explicit and implicit learning served as the primary motor learning metrics in our correlation analyses. Asymptotic measures were used to examine the overall tradeoff between the two processes at the individual level.

### Cross-Task Correlations

With respect to correlations across tasks, we had two primary hypotheses: first, that effector-independent MWM would selectively correlate with explicit motor learning, and second, that effector-specific MWM would selectively correlate with implicit motor learning. To test the first hypothesis, we computed participants’ average MWM performance in the Switch condition (collapsing across all four movements of the encoding sequence) using both variability (IQR) and average absolute angular error metrics. We then correlated these metrics with total explicit learning from the VMR task. We observed significant negative between-subject correlations between errors in effector-independent MWM and total explicit motor learning (Figure 4; variability: *r*_*pearson*_ = -0.44, *p* = 0.013; *r*_*spearman*_ = -0.46, *p* = 0.01; mean absolute angular error: *r*_*pearson*_ = -0.41, *p* = 0.024).

**Figure 4.**
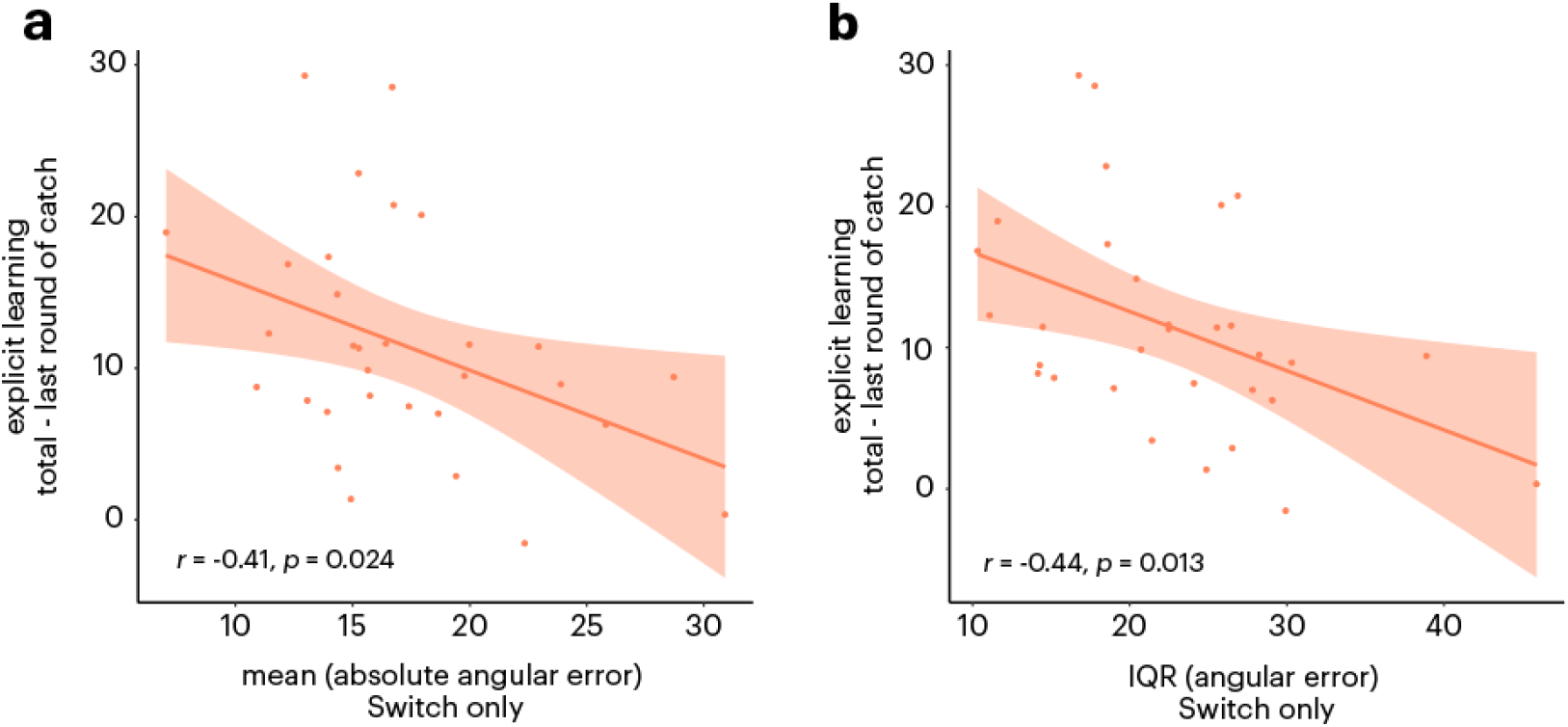
Explicit motor learning and effector-independent MWM cross-task correlations. Explicit learning from the VMR task is represented on the y-axis. The x-axis displays error in the MWM task under Switch conditions (collapsed across all sequence positions), measured as either (**a**) the average absolute angular error or (**b**) the variability of angular error.

We now turn to our second hypothesis, which concerns the predicted relationship between effector-specific MWM and implicit motor adaptation. To test this prediction, we computed effector-specific MWM in two ways: first, by measuring participants’ MWM “effector-specific benefit,” and second, by examining a recency-based “pre-washout” effect in the Same trials. We used both angular variability and absolute error metrics for completeness and correlated these metrics with total implicit VMR learning (i.e., average early aftereffects; see *Methods*).

Overall, the results supported our hypothesis (Figure 5), though they were more mixed than our findings on effector-independent MWM and explicit motor learning. For the effector-specific benefit, we observed a significant positive correlation between angular variability and implicit learning (*r*_*pearson*_ = 0.38, *p* = 0.037; *r*_*spearman*_ = 0.40, *p* = 0.025), but did not see a reliable correlation when using the average absolute angular error metric (*r*_*pearson*_ = 0.24, *p* = 0.202; *r*_*spearman*_ = 0.18, *p* = 0.323).

**Figure 5.**
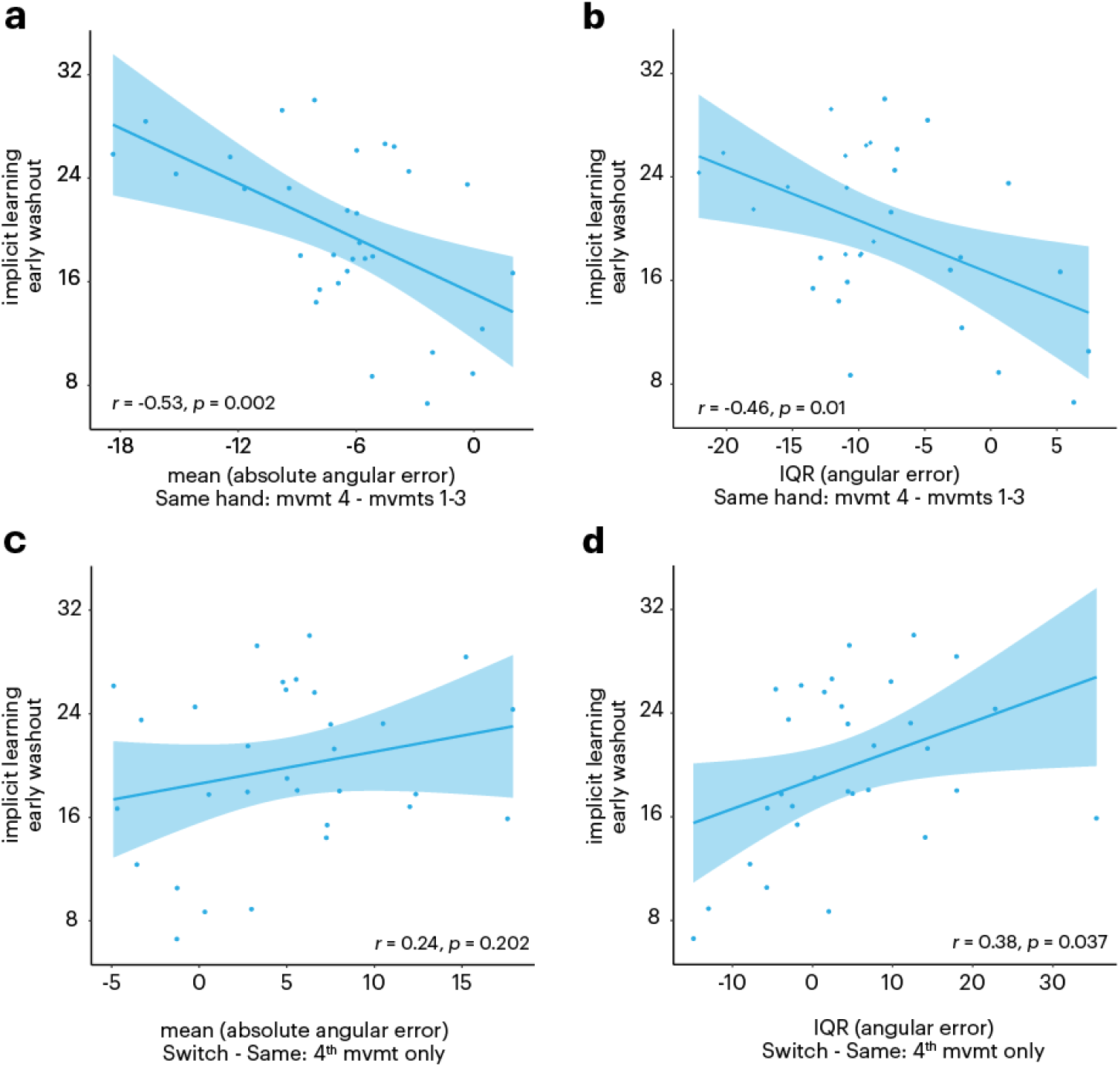
Implicit motor learning and effector-specific MWM cross-task correlations. Top: Effector-specific MWM was quantified as the difference in error between the fourth movement, which demonstrated an effector-specific benefit, and the first three movements within the Same condition. Error was measured using (**a**) the average absolute angular error and (**b**) the variability of angular error. Since this calculation is based on error, a negative correlation signifies a positive relationship between implicit learning and effector-specific MWM. Bottom: The x-axis represents the difference in error between the Switch and Same conditions for the fourth movement in the MWM task, using both (**c**) the average absolute angular error and (**d**) the variability of angular error.. This difference reflects the advantage of recalling movements with the same hand (the effector-specific benefit).

For the fourth-movement measure of effector-specific MWM, we observed a correlation between the angular variability and implicit motor learning (*r*_*pearson*_ = -0.46, *p* = 0.01; though not reliable for the Spearman metric, *r*_*spearman*_ = -0.27, *p* = 0.138), and a correlation between the average absolute angular error metric and implicit motor learning (*r*_*pearson*_ = -0.53, *p* = 0.002; *r*_*spearman*_ = -0.46, *p* = 0.009). We note that the sign change in the correlations within Figure 5 reflects the fact that effector-specific MWM in (Figure 5, bottom row) refers to a reduction in error.

It is possible that some of our correlations, especially for explicit measures, could be driven by differences in general task effort across subjects. To address this, we performed control correlation analyses using extent error metrics from the MWM task rather than angular error metrics, as the angular dimension is the only kinematic dimension that is relevant to the visuomotor learning task. If the results point to true task-specific underlying relationships between MWM and visuomotor rotation learning, the extent-error-based correlations should not be reliable. Indeed, we did not observe significant correlations between extent-based MWM error metrics and motor learning metrics (all *p*’s > 0.12).

These findings demonstrate that the fidelity of effector-independent MWM is correlated with explicit learning in VMR, and that effector-specific MWM appears to be correlated, though perhaps to a lesser extent, with implicit motor learning. Taken as a whole, the results shown in Figures 4 and 5 thus point to a double-dissociation in the relationship between MWM codes and explicit and implicit motor learning.

## Discussion

In this study, we explored the relationships between short-term memory for movements (motor working memory; MWM) and motor learning. Specifically, we asked whether the capacity to hold effector-independent and effector-specific information in MWM is selectively correlated with explicit and implicit components of motor learning, respectively. We found evidence supporting these hypotheses: (1) individuals with more robust explicit motor learning also exhibited greater memory fidelity in effector-independent MWM, and (2) individuals with a high level of implicit motor learning also exhibited greater memory fidelity in effector-specific MWM. The results add further support to the distinction between effector-specific and effector-independent representations in MWM (Hillman et al., 2024), and they offer novel insights into how MWM may contribute to long-term motor learning processes.

### Effector-independent MWM and explicit motor learning

Prior research relating working memory to motor learning has been largely restricted to examining the relationship between visual working memory and motor learning (Anguera et al., 2010; Bo & Seidler, 2009; Christou et al., 2016). Here we found that a greater contribution of explicit processes during motor learning was correlated with lower error in effector-independent MWM. Given recent findings from our lab showing no interference between visuospatial load and effector-independent MWM, these results build on previous work by identifying “motor-specific” working memory processes as potentially a distinct contributor to explicit motor learning (Hillman et al., 2024).

Our results linking effector-independent MWM and explicit motor learning are also consistent with intermanual transfer research. Transfer studies have used visuomotor and force field adaptation tasks, in which one arm undergoes adaptation and the opposite arm is tested for residual learning. Recent work has demonstrated that explicit learning is largely effector-independent (transferable across limbs) (Poh & Taylor, 2019; Schween et al., 2020), which is consistent with our results. Furthermore, a study by van Mier & Petersen (2006) found that participants who learned a maze by tracing showed significantly better transfer when tested with the opposite hand on an identical maze than on a mirrored one. This finding supports the MWM task design, where effector-independent MWM is assessed by replicating a non-mirrored movement on Switch trials.

### Effector-specific MWM and implicit motor learning

We also found evidence supporting our secondary hypothesis, that implicit motor learning may leverage effector-specific information held in working memory. We observed a significant positive correlation between implicit learning and effector-specific MWM, measured as both the memory benefit afforded by using the same hand to recall movements, and a putative recency-based “fourth-movement” metric related to recalling the most recently encoded movement with the same hand. However, when using both our absolute error and variability MWM metrics, one of these correlations – between implicit adaptation and the absolute error in the same-hand benefit metric – was not reliable (Figure 5). Thus, the correlations observed between implicit adaptation and effector-specific MWM should be taken with a grain of salt.

Our finding that effector-specific MWM correlated with implicit motor learning is broadly consistent with other findings showing that, unlike explicit motor learning, implicit motor adaptation is highly effector-specific, exhibiting minimal (if any) transfer across limbs (Joiner et al., 2013; Malfait & Ostry, 2004; Schween et al., 2020). Moreover, recent work in motor adaptation has uncovered a sub-component of implicit motor learning that is susceptible to short-term forgetting (“temporally-volatile” adaptation), biasing movements back to an unadapted state (Hadjiosif et al., 2023). These effects could relate to a possible reliance on some kind of implicit working memory cache. While our previous study found a lack of temporal decay effects in MWM (Hillman et al., 2024), those analyses were focused on an increase in memory errors over time, not a drift back to a non-adapted motor memory. Future work could attempt to more directly relate our measures of working memory and temporally-volatile implicit adaptation.

Previous work has highlighted a negative relationship between working memory recruitment and implicit motor processes (Christou et al., 2016), which may appear contrary to our results here (Figure 5). We propose that the positive relationship we observed between implicit motor learning and effector-specific MWM challenges the conventional view that any working memory recruitment impairs implicit motor learning. Instead, it suggests that interference may stem not from working memory recruitment itself, but from the specific content maintained within it.

### MWM: Eligibility traces for learning?

The correlations between effector-independent MWM and explicit learning we observed here may reflect the requirements of remembering an abstract motor plan across short timescales (Velázquez-Vargas & Taylor, 2024). This type of working memory process would be quite useful for retrieving and refining a consistent cognitive strategy during learning, and linking those strategies to observed errors during performance monitoring (Hewitson et al., 2023; McDougle, Boggess, et al., 2016; Tsay et al., 2024; Velázquez-Vargas & Taylor, 2024). In the context of implicit learning, short-term memory of movements could help link observed sensory errors to preceding movements, especially under feedback delays (Brudner et al., 2016; Kitazawa et al., 1995).

These putative functions of MWM during are akin to a kind of “eligibility trace,” a key concept in reinforcement learning (Sutton & Barto, 1998). That is, working memory can help the learner link recent actions held in memory (the “traces”) with specific observed outcomes (Curtis & Lee, 2010; Sidarta et al., 2018). This key computation could be implemented by maintaining a persistent representation of recent action kinematics. Neural recordings in monkeys performing saccades provide some direct neural evidence for a putative MWM system in the brain – these studies show substantial post-saccade working memory traces of eye movement kinematics in the dorsolateral prefrontal cortex (Calangiu et al., 2025; Tsujimoto & Sawaguchi, 2004), a key hub of the working memory system. It is plausible that these findings would generalize to other motor behaviors, like reaching. Future work could try to reveal in more detail the precise format and neural correlates of persistent working memory representations of actions that could be used to support motor learning.

### Limitations

While the current study provides evidence supporting a role for MWM in motor learning, several limitations should be considered. First, the correlational design does not allow for definitive conclusions about causal relationships. Future studies could employ experimental manipulations, such as introducing a motor working memory load, to establish a causal link between motor learning and effector-independent or effector-specific MWM. Second, our measures of explicit learning and effector-specific MWM were derived indirectly, relying on subtraction-based calculations. While these approaches are widely used, they may introduce error (Maresch et al., 2020) or confounding influences from other cognitive processes. Additionally, it is unclear how the findings from our simple lab-based task would generalize to more ecologically realistic settings. In naturalistic environments, individuals often hold information from multiple sensory modalities in working memory to guide actions—for example, vision—which could either enhance or interfere with learning. Furthermore, the relationship between short-term memory and motor learning likely varies across different age groups and skill levels. For instance, athletes and musicians may attend to or retain different types of information in MWM, influencing motor learning in distinct ways. Taken together, our findings suggest that MWM likely plays a role in multiple forms of motor learning.

## Acknowledgements

We thank Samantha Goodcase for helping collect data. We also thank the ACT Lab for helpful discussions. Work supported by grant R01 NS134754 (S.D.M.) from the National Institutes of Health.

